# The effect of data preprocessing on spike correlation analysis results

**DOI:** 10.1101/2025.11.28.691090

**Authors:** Jonas Oberste-Frielinghaus, Junji Ito, Sonja Grün

**Affiliations:** Institute for Advanced Simulation (IAS-6), Jülich Research Centre, Jülich, Germany; RWTH Aachen University, Aachen, Germany; JARA-Institute Brain Structure-Function Relationships (INM-10), Jülich Research Centre, Jülich, Germany; Theoretical Systems Neurobiology, RWTH Aachen University, Aachen, Germany

**Keywords:** spike synchrony, spike timing, artifacts, preprocessing

## Abstract

In recent years, an increasing number of large electrophysiological data sets have become publicly available, thereby providing researchers with the opportunity to analyze spike train data without conducting their own experiments. While this is undoubtedly a positive development, it increases the need for proper documentation on how the data were collected and what preprocessing was performed on the data, since interpreting analysis results in ignorance of these pieces of information can lead to wrong conclusions. An important preprocessing step is the removal of artifacts from the recordings. Electrophysiological recordings are particularly susceptible to electrical cross-talks between recording channels, resulting in artifact spikes that are coincident in multiple channels on the time scale of the data sampling rate, i.e., 1/30 ms in popular setups. The removal by signal whitening is only possible if also the raw sampled data are available, thus to eliminate this type of artifact is to remove all coincident spikes on the recording time scale to definitely avoid artifact spikes. However, given the lack of the “ground truth”, this step has the potential to eliminate, in conjunction with the artifacts, components of the data that are pertinent to the research objective. In this study, we use a modified version of the Unitary Event Analysis and demonstrate that such preprocessing results in significantly lower correlations than expected by chance even on longer time scales. We also propose a method to correct for the bias introduced by this preprocessing. Thus, slight changes in the preprocessing have potentially strong impact on analysis results and methods need to consider these effects.

## 1 Introduction

Large parts of neuroscience rely on electrophysiological recordings for the investigation of dynamical properties and computational mechanisms of the brain. The experiments that are required to obtain these data involve considerable effort and resources. Fortunately, it has become more common to share data within the community e.g., Brochier et al. (2018); Steinmetz et al. (2019); Pei et al. (2021); Chen et al. (2022), making recordings accessible for a broad range of researchers. However, there is no standardized format for reporting and each data set is highly individual. Thus researchers other than the experimenters often times do not know all information about the data even if the data is very well documented.

Typically the data undergo a number of preprocessing steps, such as frequency filtering, down-sampling, spike sorting, etc., which are not necessarily included in the description of the data. Each of these steps can influence the data and not knowing about them might impact the interpretation of analysis results, for example Bedenbaugh and Gerstein (1997); Pazienti and Grün (2006) showed how errors in spike sorting — typically a process the data user has only limited knowledge about — may affect the observed correlation structure.

Here we focus on another preprocessing step, namely the removal of artifacts. In Oberste-Frielinghaus et al. (2025), we have shown that in electrophysiological recordings correlations on very high time resolutions across multiple recording channels are ubiquitous. These are noticeable as high synchronous spike count entries in the spike population histogram of the activities of all neurons in single trials computed on the sampling resolution (here: *f*_s_ = 30 kHz), either on sorted spikes or thresholded spike data. These high entries occur much more often than expected by chance (comparison to surrogates). Correlations on this time resolution are very unlikely caused by neuronal activity and thus probably are caused by artifacts in the data. To mitigate the impact of these artifacts, previous studies either excluded whole recording channels e.g., Yu et al. (2009), potentially removing large parts of the data, or like Torre et al. (2016) removed those artifact spikes by removing any synchronous spike events at sampling rate resolution likely also including “real” spikes. In Oberste-Frielinghaus et al. (2025), we report a more elegant way for artifact removal by whitening the raw signals based on the zero-phase component analysis (Bell and Sejnowski (1997)). However, since this cleaning method needs to be applied to the raw continuous data, it is not an option for data users who do not have access to the raw data.

In this work, we illustrate the effect of data cleaning (artifact removal) as done in Torre et al. (2016). In particular we study the effect of this artifact removal on the results of spike correlation analysis on the millisecond time scale, using the Population Unitary Event analysis (UE_pop_), newly introduced here. We choose this type of analysis because it uses many spike trains in parallel, and thereby the effect of errors is more apparent due to the large amount of data. For the demonstration of the effect of the artifact removal, we apply the UE_pop_ to simulated data in which we emulated artifacts as well neuronal correlations as observed in neural recordings. Since we know the ground-truth processes underlying these data sets, they are ideal to exemplify the effect of the artifact removal under various different constellations. We find that this removal method leads to an increase in false negative results. Additionally, we suggest a correction for UE_pop_ to account for the effects of the artifact removal and to avoid these false results.

## 2 Methods

### 2.1 Recording and analysis time scales

We assume the raw data to be recorded at 1/30 ms resolution (30 kHz sampling rate), as usually nowadays performed in typical electrophysiological recordings. These data are typically high pass filtered (above ∼ 500 Hz) to get the spiking activity, which are then spike sorted to get single unit spiking activities at that high time resolution. Figure 1a, top sketches such single units spike data of *N* simultaneously recorded neurons. To apply data analysis on such data we apply a temporal binning with e.g. 1 or 5 ms bin width. If more than one spike are in a bin, we clip them to an entry of 1 (see Figure 1a, bottom) for the intended subsequent Population Unitary Event analysis (UE_pop_), described in the following section.

**Fig. 1.**
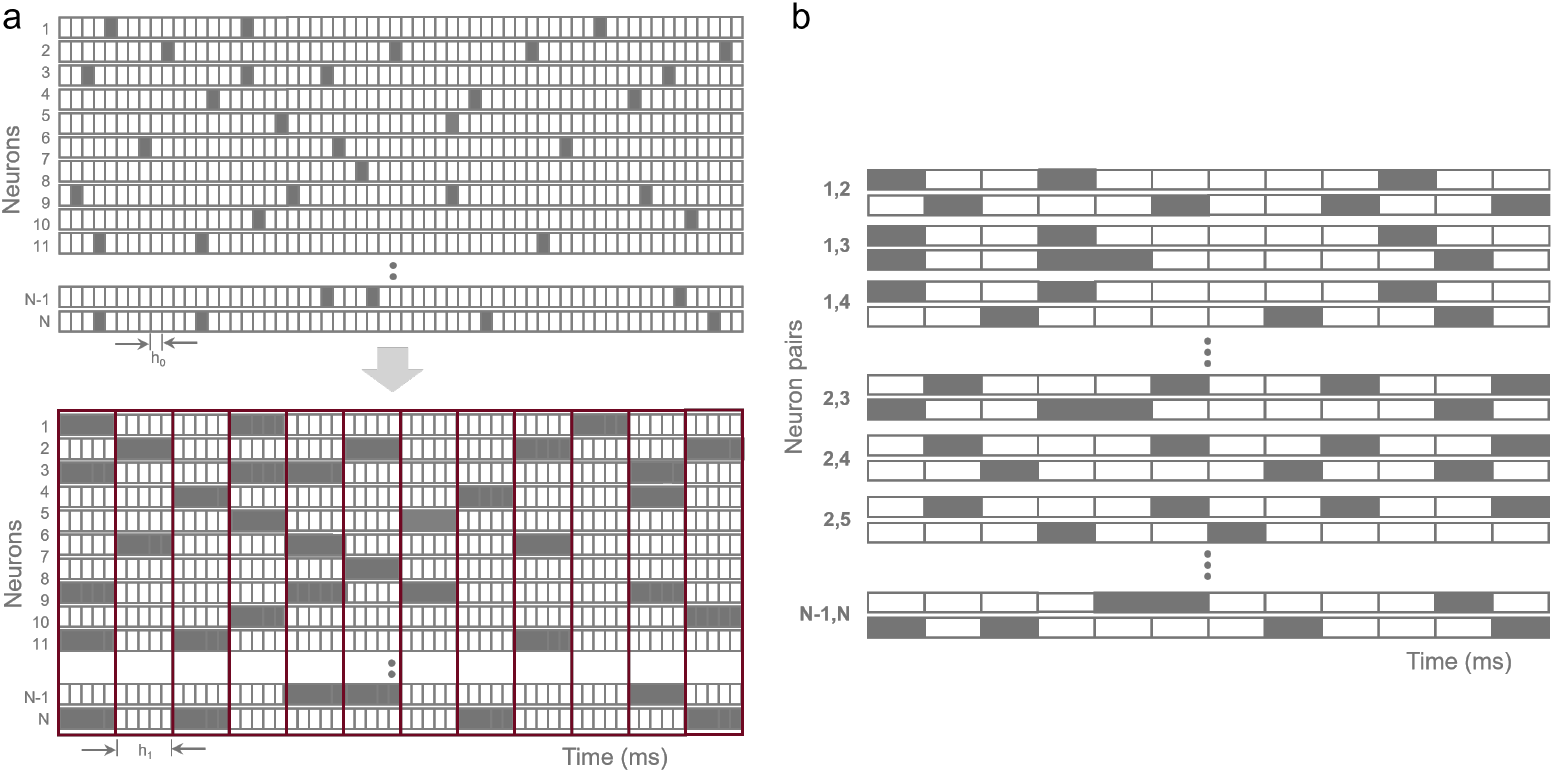
Sketch of spike data on original time resolution to analysis time resolution. (a) The top panel shows the N spike data (spikes indicated by filled bins) on the *h*_0_ = 1*/*30 ms time resolution. Below are the same data but now after binning to*h*_1_ = 1 ms bins. (b) Sketch of the UE_pop_ analysis method. All pairs of binned spike data (no repetition of identical pairs) are illustrated. From each of those pairs (*i, j*) the empirical number of synchronous events 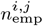 and the expected number of synchronous events 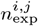 are derived. After summing the empirical number of synchronous spike events into 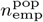 and the expected number of spike events 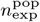 the p-value and the surprise are computed from these two values. The result of this evaluation is represented in the middle of the analysis time window of *T*_0_ ms and performed in sliding window manner.

### 2.2 Population Unitary Events Analysis

The goal of UE_pop_is to derive if there exist excess synchronous spike events in data of say *N* = 100 parallel spike trains. For this we first decide on a time resolution at which spike synchrony is considered, here *h*_1_ = 1-5 ms, and reduce the time resolution by binning the data to that time scale. The contents per bin is 1 if there are 1 or more spikes in a bin, or 0 otherwise. Thereby we end up with binary data. After this we apply UE_pop_to the data, which is an extension of the Unitary Event analysis (UE) (Grün et al. (2002a,b)) to many pairs of spike trains. To account for nonstationary firing rates in time, we apply - as for UE - the analysis in a sliding window manner of e.g. a width of *T*_0_ ms (typically 100 ms). Then we consider all pairs of neurons (no duplication, i.e., *N* (*N* − 1)/2 pairs), extract from each neuron pair (*i, j*) in that time window the empirical number 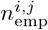 of synchronous events (just by counting), and then sum them across all pairs: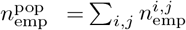. Similarly we derive the expected number 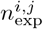 of synchronous spike events pair by pair and sum them across all pairs: 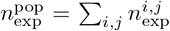. The expected number for a pair of neurons is computed on the basis of each of their spike counts *c*_*i*_ within the window to derive the probability of bin occupation by dividing the spike count by the number of bins *M*_1_ = *T*_0_/*h*_1_: *p*_*i*_ = *c*_*i*_/*M*_1_. The expected number for a neuron pair (*i, j*) is then derived as their joint probability under independence multiplied by the number of bins in 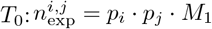.

The significance of potential excess synchrony is tested with a p-value *p*_*v*_ assuming the empirical synchronous events Poisson distributed, which is the case for Poisson spike trains. For spike trains that deviate from Poisson, we suggested to use surrogate methods (Louis et al. (2010); Stella et al. (2022)), which could also be applied here, but for the sake of simplicity we illustrate the impact of preprocessing on simulated Poisson data. The p-value is computed as the sum of a Poisson distribution of mean 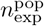 from 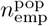 to infinity, which provides the probability to get 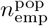 number of synchronous events or even more (Palm (1981)). From the p-value we derive the surprise measure *S*(*p*_*v*_) = log((1 − *p*_*v*_)/*p*_*v*_), a measure that fluctuates around 0 if the data are independent, is negative for a lack of synchrony, or is positive if excess synchrony exists. If *p*_*v*_ is smaller than a predefined significance level (conventionally 5% or 1%), we consider the empirical occurrences as significant on the respective significance level.

### 2.3 Simulating ground truth data

For the demonstration of the effect of preprocessing on UE_pop_, here the artifact removal, we use simulated data where we have all parameters under control (“ground truth data”). The simulations are done on the basis of a stochastic point process, the Compound Poisson Process (CPP) (Staude et al. (2010)). For all types of data we will use, a) independent Poisson processes, b) artifact data, c) neuronal correlations. For the latter two we use a CPP with different parameters, and depending on the data needed, we merge the resulting data respectively. To adjust the processes to artifact observations in experimental data, we use as a reference for the degree of artifacts the measurement of the complexity distribution, introduced in the following.

#### 2.3.1 Population spike counts and complexity distribution

From parallel processes (or measurements) binned at *h*_0_ time resolution, we form the population histogram *Z*(*s*), i.e., the sum of spike events per bin across all *N* processes 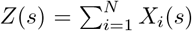, where *X*_*i*_(*s*) is the spike count of the i-th neuron in the s-th bin, i.e., the time interval [*sh*_0_, (*s* +1)*h*_0_]. The variable *Z*(*s*) defines the complexity of the population spike count at the s-th bin, irrespective of the neurons that emit these spikes. We form the complexity distribution (Grün et al. (2008); Staude et al. (2010)) as the probability distribution of the complexities Z(s): *f*_*Z*_(*l*):= *Pr*(*Z* = *l*).

In Oberste-Frielinghaus et al. (2025) and Torre et al. (2016), we used the population histogram Z(s) (on the recording time scale *h*_0_) and the resulting complexity distribution *f*_*Z*_ to identify artifacts. To do so we compared the complexity distribution of surrogate data generated from the original data, i.e., randomized versions of the original data to make the spike trains independent (e.g. the spike train shift method, (Stella et al. (2022))) to the complexity distribution of the original data. The complexity distribution of the surrogate data lacked higher complexities which were observed in the original data. Below in Section 3.1 we make use of the complexity distribution obtained from experimental data for the generation of simulated spike trains containing artifacts (Figure 2d).

**Fig. 2.**
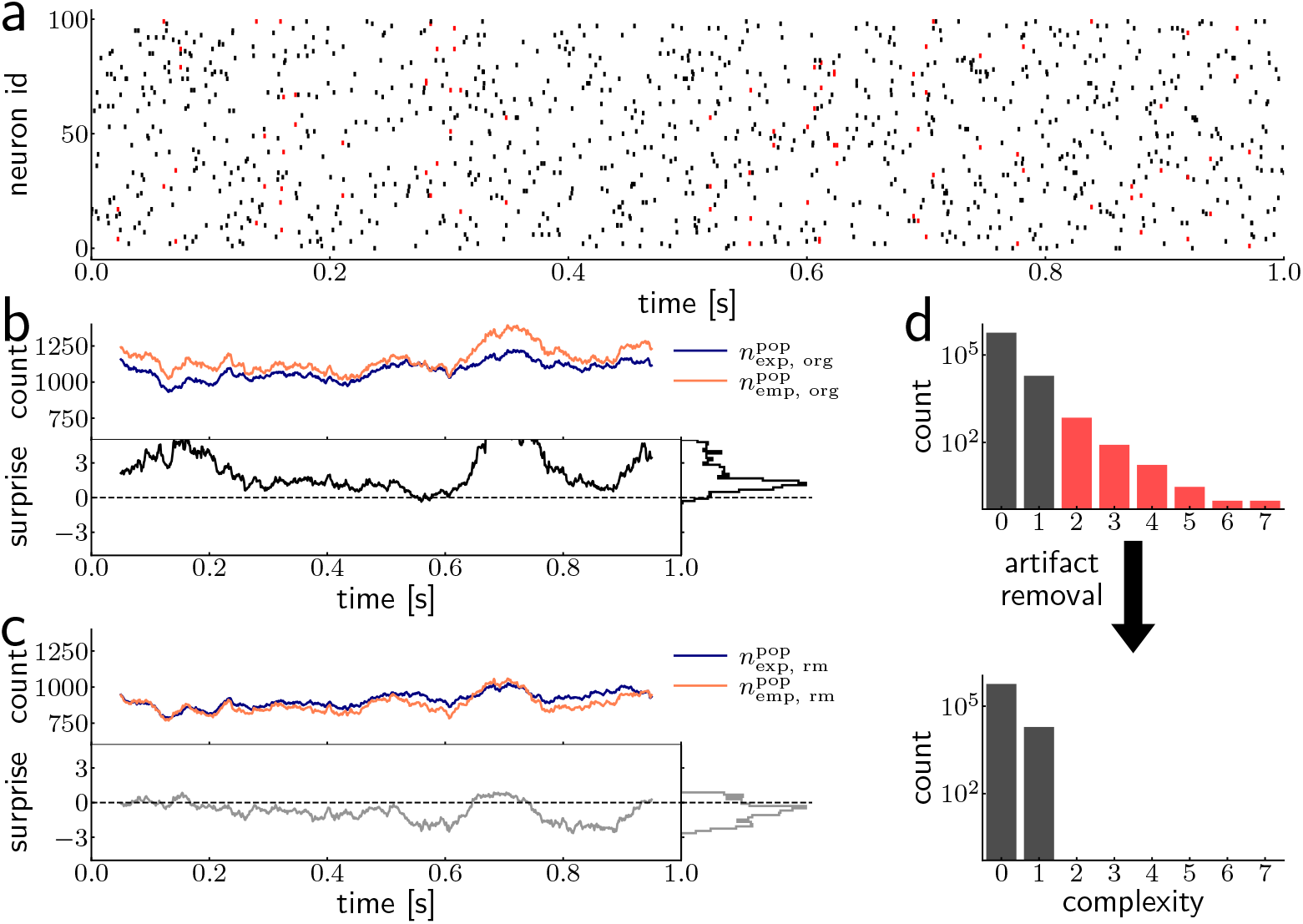
Effect of artifact removal. (a) Raster plot of parallel spike trains (*N* = 100) for one trial over 1 s containing background firing and artifacts, simulated using CPPs at a temporal resolution of *h*_0_ = 1*/*30 ms. The background rate of each spike train is 10 Hz (black dots). The amplitude distribution is set identical to the complexity distribution obtained from an experimental dataset (d), which is known to contain synchronous artifacts (Oberste-Frielinghaus et al. (2025)). The red dots indicate artifact coincident spikes on the 1*/*30 ms time scale, which are to be removed as artifacts. (b) Result of the sliding window UE_pop_ performed on the binned and clipped (at *h*_1_ = 1 ms resolution) data. The top panel shows the time course of the expected and empirical number of spike coincidences (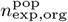 (dark blue) and 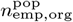 (orange)). Below is the time course of the surprise (black), with its marginal distribution to the right. The distribution of the surprise shows a mean of about 2, indicating excess synchrony, almost throughout. (c) The same measurements (top: 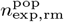 (dark blue) and 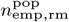 (orange)) and the surprise below) of the same data after the artifact removal. Here the surprise distribution is almost completely below 0, which reflects that the empirical number (orange) in the data after the removal is mostly below the expected number (dark blue). (d) The complexity distribution of the spike coincidences at the *h*_0_ time resolution (top) and below on the same time scale after the artifact removal. In the top histogram the red entries mark complexity entries > 1, which are removed as artifacts, corresponding to the red dots in panel a.

#### 2.3.2 Compound Poisson Process

To generate parallel point processes that contain correlations between the processes we make us of the Compound Poisson Process (CPP) as introduced by Staude et al. (2010). We will make use of this type of process for simulating parallel spike trains that contain artifacts or neuronal correlations, which will be explained in detail below. The idea is the following: a hidden point process *z*(*t*), called carrier process, is generated with a particular firing rate *α* in continuous time. In addition, we specify an amplitude distribution *f*_A_(*a*) which defines a probability distribution from which integer-valued ‘amplitudes’ *a* are drawn with their respective probabilities. To construct the *N* parallel spike trains containing correlations, amplitudes *a*_*j*_ are drawn from *f*_A_ for all events *t*_*j*_ in the carrier process, resulting in *z*(*t*) = ∑_*j*_ *δ*(*t*−*t*_*j*_)·*a*_*j*_. The individual “child” processes *x*_*i*_(*t*) (*i* = 1, 2, …, *N*) are then constructed by copying every event at *t*_*j*_ of the carrier process into *a*_*j*_ randomly drawn child processes at the same time as *t*_*j*_. Thus, the probability for amplitude *a* = 1 defines the probability for complexity *ξ* = 1, amplitude *a* = 2 for complexity *ξ* = 2, and so on. This way, correlations of order *ξ* are induced in the child processes whenever events in the carrier process *t*_*j*_ are copied into *ξ* = *a*_*j*_ child processes. The firing rates of the child processes are derived as the firing rate *α* of the carrier process divided by the number of child processes. The *N* child processes retain the same time resolution as the carrier process. In our context we then discretize the resulting processes defined on continuous time to the time resolution of the sampling rate, i.e., *h*_0_ = 1*/*30 ms and binarize the content to 0 and 1.

#### 2.3.3 Independent parallel spike trains

As independent spike trains we generate *N* = 100 realizations of stationary, independent Poisson processes on continuous time of rate *λ*_*b*_ =10 sp/s and of a duration of *T*_0_ = 100 ms and across multiple trials. These data are then discretized to the time resolution of *h*_0_ = 1/30 ms and binarized to 0-1 contents.

#### 2.3.4 Parallel spike trains with artifacts

To generate parallel spike trains that just contain artifacts as identified by a complexity distribution of experimental data measured at the sampling resolution (1/30ms). Therefore we generate a CPP and use the measured complexity distribution and the amplitude distribution, however with an entry of 0 at *ξ* = 0 and 1, since the independent background activity is generated independently (Section 2.3.3). We estimated from experimental data that the artifact spikes had a rate of *λ*_*a*_ = 0.5 sp/s such that the carrier rate resulted in *α* = *N* · *λ*_*a*_ = 50 sp/s. Also here, the resulting data are then discretized to the time resolution of *h*_0_ = 1*/*30 ms and binarized to 0-1 contents.

These resulting data are then merged (on *h*_0_) with the independent data generated through the independent Poisson processes (Section 2.3.3) as uncorrelated background and the artifact data generated here. The resulting data are shown in the raster plot in Figure 2a, the artifact spikes are shown in red. The resulting complexity distribution (on *h*_0_) Figure 2d includes the background activities (black) and the artifacts with complexities 2-7 (red) without mentioning that some background spikes may also generate - with low probability - by chance also pair entries.

#### 2.3.5 Parallel spike trains that include ‘neuronal’ correlations

Physiological spike correlations, i.e., correlations that are assumed to originate from neuronal spike correlations, are generated by another CPP. Here we introduce just pairwise correlations, i.e., by use of an amplitude distribution of *f*_A_(*a*) = 1 for *a* = 2 and 0 otherwise, with a correlation rate *λ*_c_ = 1sp/s (as found in experimental data (Maldonado et al. (2008))). Thus the carrier rate is *α* = 100 · *λ*_c_. For each spike of the carrier process two neurons are drawn randomly into which a spike is copied. Thus each correlated spike of one neuron is correlated with another neuron.

Since neuronal correlations are typically less precise than 1/30 ms, but rather on a 1-5ms scale (Riehle et al. (2000)), we jitter each carrier spike before insertion into the child process by a random amount *δ*_*j*_, uniformly drawn from an interval [−Δ,Δ]. As a result the spike coincidences are synchronous up to a precision of ± Δ,thus 2 ·Δ. Then the processes are discretized and binarized to *h*_0_ = 1*/*30 ms.

Finally, the processes containing neuronal correlations are merged with independent processes that have a reduced rate for each neuron, *λ*_uncorr_ = *λ*_*b*_ − *λ*_c_, such that the firing rate of the spike trains after introduction of the correlations equals to *λ* = 10 sp/s.

#### 2.3.6 Parallel spike trains that include artifacts and ‘neuronal’ correlations

For validation of the correction method for the errors caused by artifact removal in presence of neuronal correlation, we also generate ground truth data sets that include artifact spikes and correlated neuronal spikes. To do so we first generate independent background spikes (see Section 2.3.3), then spike trains that contain artifacts (see Section 2.3.4), and also spike trains with neuronal correlations and merge the binarized versions (on *h*_0_ resolution) into one data set.

## 3 Results

### 3.1 Removal of artifacts leads to decreased surprise

Figure 2 illustrates the effect of artifact removal on the UE_pop_ analysis. The data shown in Figure 2a are *N* = 100 parallel spike trains containing artifacts, generated by a CPP (Section 2.3.4) at the time resolution *h*_0_ = 1*/*30 ms. The amplitude distribution *f*_*A*_ was chosen as the complexity distribution found in an experimental dataset, which is known to contain synchronous artifacts (Oberste-Frielinghaus et al. (2025)). The background firing rate of each the processes was chosen as 10 sp/s (by Section 2.3.3). Thus the complexity distribution in Figure 2d illustrates the combination of the background activity (black) and the artifacts (red).

The spike data shown in Figure 2a were then analyzed by UE_pop_ performed at *h*_1_ = 1 ms time resolution in a sliding window manner (*T*_0_ = 100 ms). In Figure 2b we can observe the resulting empirical 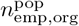 (orange)) and the expected number 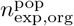 (dark blue) of spike synchrony, and below the resulting surprise. The fluctuations of the empirical and expected numbers in time result from stochastic fluctuations of the generated stationary processes. Over time the empirical number is above the expected, with a slightly fluctuating difference between the two, resulting in a modulating surprise with values almost always above 0 with a mean at around 2. Thus most of the time the surprise shows excess synchrony (above p-value = 0.05, i.e., a surprise of 1.67). However, this excess synchrony is not due to neuronal correlation but due to the artifacts contained in the data.

In Figure 2c we see the result of the UE_pop_ after removal of the artifacts, i.e., the removal of all coincident spikes at the time resolution of *h*_0_ = 1*/*30 ms, as indicated in Figure 2d by the complexity distribution after artifact removal (bottom). Now the relation of the empirical (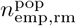, orange)and the expected number(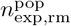, dark blue) are nearly reversed: the expected number of coincidences is almost always above the empirical number of coincidences, which leads to surprise values almost always below a surprise of 0, with a mean of about −1, thus negative correlations.

This results show us two things: 1) artifacts can lead to artificial excess correlation in uncorrelated, independent neuronal data and 2) the approach to remove the artifacts based on the removal of all synchronous events at the recording time resolution *h*_0_ overcompensates. If data were independent and not correlated, the surprise should not be overall negative, but should fluctuate around 0. The conclusion is that we removed too much synchronous spikes, likely chance coincidences on the *h*_0_ time scale. This is further studied in the following.

### 3.2 Removal of highly synchronous spikes leads to decreased surprise in independent data

To observe the effect of the artifact removal more clearly, we now deal with data that contain no artifacts (i.e., independent spike trains), and again apply the preprocessing step of removing all coincident spikes on the level of the sampling rate resolution *h*_0_ as before. Since this only concerns the spike timing at a very high resolution, i.e., 1/30 ms, only a very small amount of synchronous spikes is removed. In our simulated data, we have 20 trials of 1 s duration in which 100 neurons are generated, each neuron simulated as a Poisson process of a rate of 10 sp/s. In Figure 3a, we see the resulting spike times as a raster plot for one example trial under these conditions. If we now apply the suggested preprocessing, we remove all (synchronous) spikes marked in red. In terms of the complexity distribution, this refers to all spikes with a complexity equal to or larger than two (red bars in Figure 3b). Since these are all chance synchronous spike events at this high time resolution, those are rare.

**Fig. 3.**
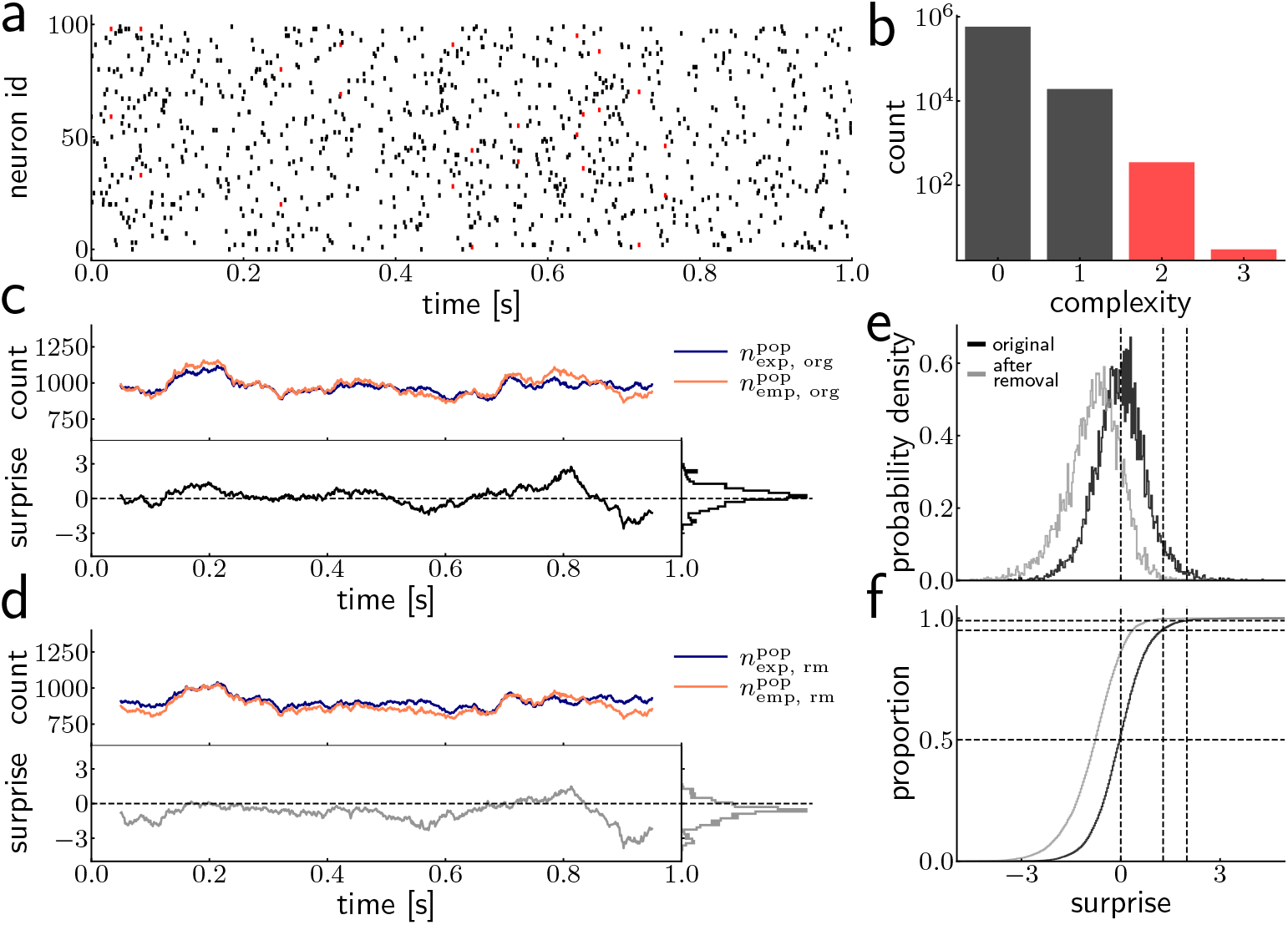
Effect of removal of precise coincidences on the UE_pop_. (a) Example raster plot for one trial over 1 s of 100 neurons in parallel with each originating from a CPP that generates independent Poisson processes of 10 sp/s (Section 2.3.3). The dots correspond to the spike times of the neurons. Red dots mark spikes with a complexity of 2 or larger, black dots mark spikes with complexity 1. (b) Complexity distribution of all spikes over 20 trials, analogous to Figure 2d. (c) Top row, result for 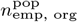 and 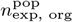 calculated by a sliding analysis with a window of 100 ms and 20 trials of the same setup as described in a). Bottom row, the resulting surprise from these values with the marginal distribution of the surprises on the right. (d) Same as c) but applied to the data after removing all spikes with a complexity of 2 or larger - as if there were artifacts in the data. Here the surprise is reduced to a mean below 0. (e) Distribution of the surprise values 1000 data sets, each consisting of 100 neurons with each a rate of 10 sp/s, recorded over 100 ms for 20 trials (without removed spike coincidences, black) and d) (with removed coincidences, gray). The vertical lines indicate 50%, the 5% and the 1% significance thresholds for the surprise (left to right). (f) Cumulatives of the distributions shown in e). The horizontal lines mark the proportion of 50%, 95% and 99% of the data sets (bottom to top).

Applying the sliding window UE_pop_ to this dataset without preprocessing, 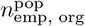 and 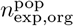 roughly match each other and the resulting surprise fluctuates around zero (Figure 3c), which is expected from independent spike trains. In comparison, after the preprocessing, the surprise is shifted to negative values (Figure 3d), i.e., we find slightly less coincidences as we expect from independent data. Thus, the removal of the synchronous events as suggested as a preprocessing step, however, has here a notable impact on the analysis result. Figure 3e shows the probability density function (PDF) of the surprise per time window before (black) and after (gray) the preprocessing, estimated from 1000 simulations of independent spike train datasets. The preprocessing shifts the obtained surprise to negative values (gray) as compared to the results without preprocessing (black), which is also visible in the cumulative distributions of the two PDFs (Figure 3f). For the independent data, the proportion of time windows below a given surprise should match the underlying null-hypothesis, in concrete, 95% of the analysis windows lie below the significance threshold of 5% and mutatis mutandis for a significance threshold of 1%. As we see in Figure 3f, this is the case for the data before preprocessing (black), whereas in the preprocessed data a substantially fewer number of the analysis windows are detected as significant (gray).

We conclude from this analysis that the preprocessing step of artifact removal, i.e., the removal of all coincident spike events at the high time resolution *h*_0_, also removes chance coincidences. This could lead, in the analysis of correlated spike trains, to a potential underestimation of existing neuronal synchrony which will be explored in the following. ^1^

### 3.3 Preprocessing removes more coincidences than expected by chance

As shown in the previous section, the preprocessing step intended to mitigate a problem leads to a large change in the result we get from the UE_pop_. Surprisingly, a small change of the data on the sampling resolution, here the removal of chance coincidences only, impacts the result of the UE_pop_ that operates on a coarser time scale. Through the preprocessing on the *h*_0_ time scale, roughly 50-100 spikes are removed within one analysis window per dataset (duration: 100 ms, 20 trials) (Figure 4a). This corresponds to 5% or less of the spikes, which leads to a reduction of the expected coincidence count by ∼ 10% or 100 coincidences (Figure 4b). At the same time, the empirical coincidence count is reduced slightly more (approx. 150 coincidences) (Figure 4c), which leads to the impactful result that the resulting surprise is below zero (as seen in Figure 3) and thus indicates negative or lacking correlations. We conclude that the expected coincidence counts are less reduced by the removal of coincidences at *h*_0_ than the empirical counts. As a consequence, the reduction of the expected coincidence counts after the preprocessing is not sufficient to account for the reduction of the empirical coincidence counts caused by the preprocessing. This led us to a correction method introduced in the following.

**Fig. 4.**
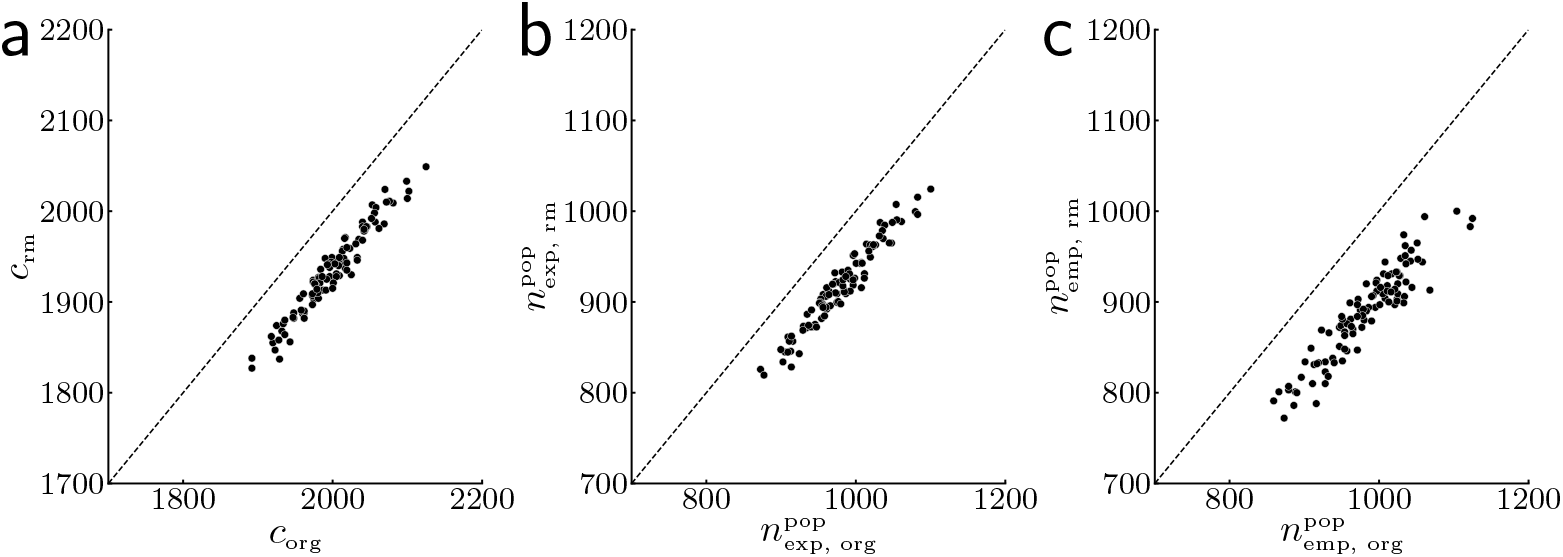
Impact of removal of synchronous spike events on *h*_0_ on the observed parameters on *h*_1_. (a) Spike count of all neurons (*N* = 100) in an analysis window of 100 ms duration over 20 trials, before (*c*_org_) and after (*c*_rm_) artifact removal. Each point corresponds to one data set, i.e., one simulation with the same parameters. (b) Expected coincidence counts for the same data sets before (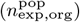) and after removal (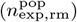). (c) Empirical coincidence counts for the same data sets before (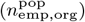) and after removal (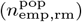). All counts are derived at a time resolution of *h*_1_.

### 3.4 Correction method for the overcompensation by the artifact removal

The results obtained so far lead to the conclusion that the artifact removal by mistake also removes chance coincidences of independent spikes occurring on the *h*_0_ time scale (i.e., data sampling time scale), which should have been kept in the empirical coincidence counts. This removal of chance coincidences causes the empirical count to systematically fall behind the expected coincidence count on the *h*_1_ time scale. The latter also include the expected chance coincidences on *h*_0_. Based on this insight, we next derive a correction of the expected coincidence count by estimating how many chance coincidences on *h*_0_we lose due to the artifact removal.

We consider independent spike trains of duration *T*, with *M*_0_ bins at *h*_0_ (e.g. 1/30 ms), leading to *M*_0_ = *T*/*h*_0_ bins. As a first step, we aim to derive, based only on the spike counts on *h*_0_ after artifact removal,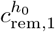 and 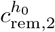 of neuron 1 and 2, respectively. Their respective firing probabilities 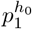 and 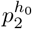 (on *h*_0_) before artifact removal are, according to the setup above,

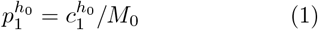

and

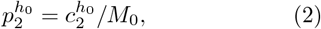

where 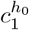 and 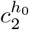 are the (unknown) spike counts of neuron 1 and 2, respectively, before artifact removal. Given these probabilities, the expected number 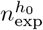 of chance spike coincidences on *h*_0_ within the analysis window *T* is obtained as

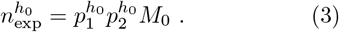

Since the spikes constituting these coincidences are the ones removed by the artifact removal, the following relations between the spike counts before and after the artifact removal hold:

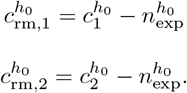

Dividing both sides of these equations by *M*_0_ and applying the relations (Equation 1), (Equation 2) and (Equation 3), we get:

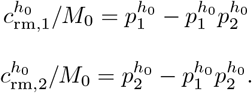

We can solve these equations for 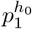 and 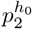 to obtain these probabilities expressed in terms of 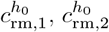 and *M*_0_ as follows^2^:

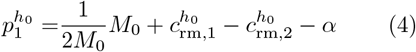

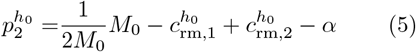

With 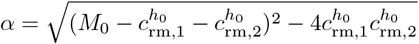.

Thus, (estimates of) the firing probabilities before the artifact removal can be derived from the spike counts after the artifact removal.

Recapitulating the UE_pop_ analysis, excess spike synchrony between a pair of neurons is detected based on a comparison between empirical and expected number (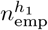 and 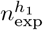, respectively) of spike coincidences on the time scale *h*_1_ within an analysis time window *T*, with *M*_1_ = *T*/*h*_1_ bins of width *h*_1_. The expected number is estimated based on the spike counts of the respective neurons within the analysis window, counted on *h*_1_ with clipping. With these spike counts 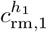 and 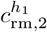 for neuron 1 and 2, respectively, the expected coincidence count 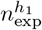 is obtained as 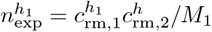.

As shown in Figure 4, after the artifact removal, the expected number 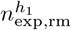 of spike coincidences is consistently higher than the empirical coincidence count 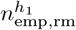. This is because 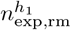 can only represent the expected coincidence count for the case where 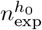 spikes are *independently* removed from each of the two spike trains, while the artifact removal selectively removes 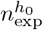 coincidences from both spike trains, and hence reduces 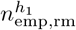 more than by an independent removal of 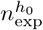 spikes. Given this consideration, we correct 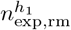 for this excess removal of coincidences by subtracting the number of excessively removed coincidences. Thus we reach the following correction of 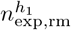:

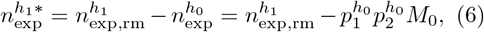

where the relation (Equation 3) is used between the middle and the right-hand side of the equation, and 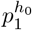 and 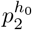 should be derived from 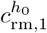 and 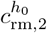 as described above (Equation 5). Note that for the corrected expected, corrected count 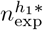 we implicitly assumed that there are no multiple coincidences on *h*_0_ within an *h*_1_ bin, and that all the 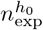 coincidences on *h*_0_ do not coincide with coincidences on *h*_1_, or in other words there is no clipping of artifacts.

We test the performance of this correction method first on independent spike train data. The black and gray curves in Figure 5 are the same as those in Figure 3, i.e., the cumulative distributions of the surprise values from UE_pop_ analysis applied to the original independent data (black) and after artifact removal (gray). The shift of the gray curve to the left reflects the overestimation of chance coincidences by the uncorrected expected coincidence count 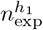 as explained above. The dashed green curve in Figure 5 represents the cumulative surprise distribution from the UE_pop_ analysis applied again to the data after artifact removal, but using the corrected expected coincidence count 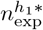 (see Equation 6). The correction works as expected: the cumulative distribution curve (dashed green) matches the curve for the original independent data (black) almost perfectly. Thus, the proposed method properly corrects the expected coincidence count in the case of the artifact removal on independent spike train data, restoring the correct analysis results expected from independent data.

**Fig. 5.**
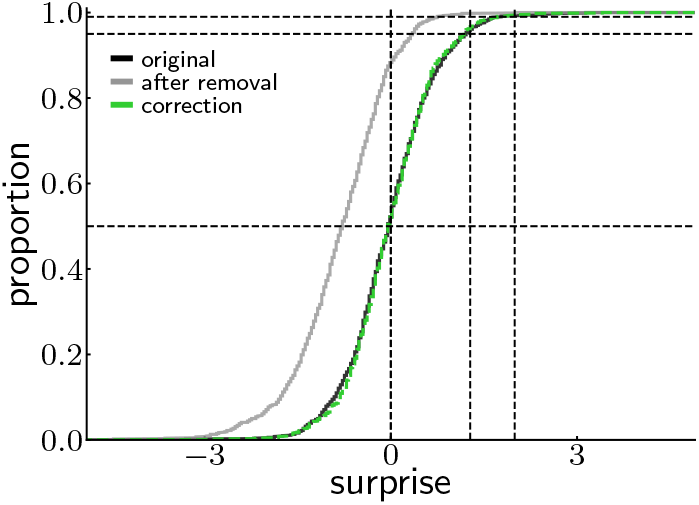
Correction of reduced correlations by artifacts removal. The graphs in this figure show the cumulative probability distributions of the surprises from the UE_pop_ for 1000 generated data sets as independent pro- cesses (see Section 2.3.3). The parameters are the same as used in Figure 3 (N=100, rate=10sp/s, trials=20, duration=100ms). Black: Original data without any removal of chance synchrony on the 1/30 ms resolution; red: after removal of chance coincidences; green: after correction.

### 3.5 Impact of the removal on correlated data

Until now, we have only investigated the removal of artifacts in the case where the underlying processes were independent. Here we apply the correction to data containing different types of spike correlations: neuronal correlations of different temporal jitters, and then also with artifacts. This allows us to validate our analytical derivations also for cases in which the data are actually correlated or contain artifacts. Correlated spikes are also generated by a CPP (see Section 2.3.5) with a temporal jitter *δ*_*j*_ (− Δ ≤ *δ*_*j*_ ≤ Δ) that mimics what we typically find in experimental, neuronal spike trains (±1-5 ms). The correlated spikes generated in such manners are then injected into background activity, i.e., independent spike trains (see Section 2.3.3). Our particular interest here is to study the effect of the proposed correction method on the performance of UE_pop_ in detecting coincidences of correlated spikes, and how these depend on the width Δ of the temporal jitter.

To this end, we first examine the effect of the correction on data containing only physiologically plausible spike correlations, without artifacts. Figure 6a left shows the results of the UE_pop_ on correlated data with a jitter width of Δ = ± 1 ms. The cumulative distributions of the surprise are obtained with the uncorrected expected coincidence count 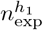 from the original (black) and the artifact removal preprocessing step (although there are none) (gray) data, and from data where artifacts were removed and afterwards the correction was applied, i.e., the corrected expected coincidence count 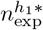 (dashed green). The difference here from the case composed of mere independent data (Figure 5) is that all three curves are shifted to the right, reflecting the spike correlation introduced into the data. Despite this shift, the corrected surprise distribution (dashed green) is relative close to the original distribution (black). In sum, these results demonstrate that the coincidence detection power of the UE_pop_, which is degraded by the artifact removal, is fully restored by using the correction of the expected coincidence count.

**Fig. 6.**
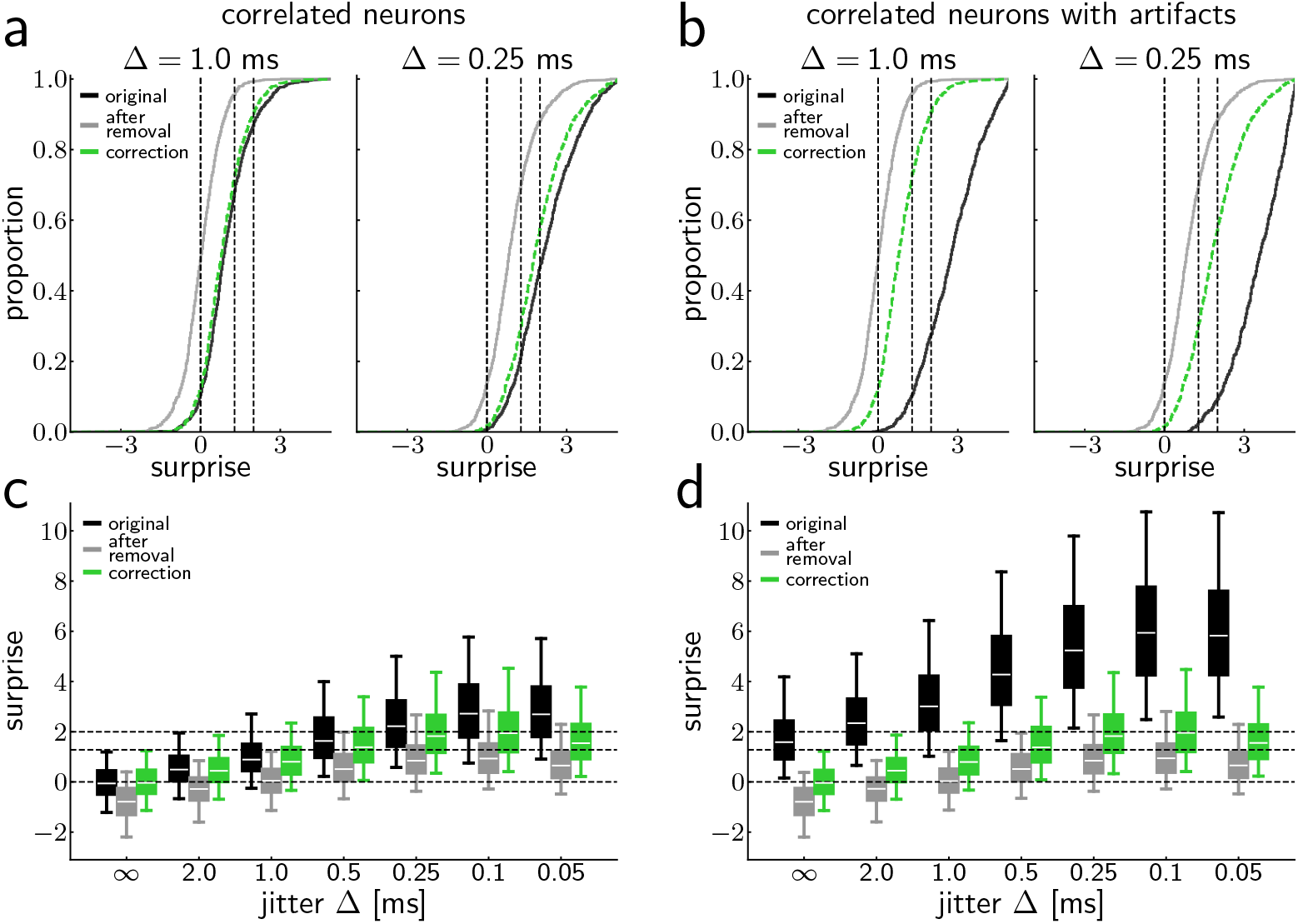
Detectability of neuronal correlations after removal of artifacts. (a) Cumulative probability distributions of the surprises from the UE_pop_ (on *h*_1_ = 1ms) for 1000 generated data sets generated as correlated processes without artifacts (see Section 2.3.6, N=100, rate=10sp/s, trials=20, duration=100ms, *r*_*c*_ = 1 sp/s into one neuron pair) with jitter Δ = 1 ms (left) and Δ = 0.25 ms (right) for the original data (black), the data after artifact removal (grey) and the corrected result (green). (b) Same as in a, but in each of the data sets also artifacts were inserted. (c) Box plots for the surprise distributions for different jitters from Δ = ∞ to 0.05 ms (Δ = ∞ refers to uncorrelated processes). The white line marks the median, the box spans the range between the first (25%) and the third quantile (75%), the whiskers range from the box to the 5th percentile and 95th percentile. The dashed lines indicate a surprise of 0, at the 5% level, and the 1% level. (d) Same as c, but for data that included artifacts.

Next, we examine whether the correction works equally well on data containing correlation with a narrower or wider jitter width than 1 ms. Figure 6a, right shows the same graphs as shown left, but obtained from the correlated data with a jitter width of 0.5 ms, i.e., the spike coincidences are temporally partly more precise than in the previous case. Due to this enhanced coincidence precision, the cumulative distribution curves shift further to the right, since more coincidences are captured within the 1 ms analysis bin (used here) as they become more precise. A rough approximation of the amount of jittered coincidences within the bin of *h*_1_ = 1 ms to estimate, how many bins of *h*_0_ = 1/30 ms of the total jitter (2 · Δ) fall into the 30 · 1/30 ms bins within *h*_1_ = 1 ms, thereby neglecting the cutting of coincidences across multiple bins of 1ms. Thus a jitter width of 0.5ms covers in total 1ms, such that almost all coincidences should be in *h*_1_ = 1 ms (neglecting the fission of coincidences by disjunct binning).

Thus in the case of a jitter of 1ms less coincidences fall into a bin *h*_1_, which explains the shift of the surprise curves to the right for a jitter of 0.5ms. Regarding the effect of the correction, the corrected surprise distribution (dashed green) resembles the original distribution (black) again, but in this case a noticeable gap can be seen between the two. Thus, the effect of the correction appears to depend on the size of the jitter of the correlated spikes or, in other words, the precision of spike coincidences of physiologically plausible spike time correlation. This leads us to systematically scan a range of jitter widths to compute the respective surprise distributions, and examine how their relative positions along the surprise axis change. Figure 6c shows box-and-whisker plots of the surprise distributions obtained from correlated data with different jitter widths Δ (x-axis), ranging from ∞, 2 to 0.05 ms, sampled approximately logarithmically. The surprise distributions generally shift towards larger surprise values as the jitter width decreases, because by decreasing the jitter, more and more correlated spikes fall into the *h*_1_ bin. This is the case for the surprise distribution from the original correlated data before artifact removal (black) over the entire range of the jitter widths examined here. In comparison, the distributions of the data after artifact removal (gray) and corrected (dashed green) show an increasing drop in surprise with decreasing jitter widths. This happens because the likelihood of an exact coincidence between correlated spikes increases as the jitter width decreases, resulting in more coincident spikes on *h*_0_ being removed by the artifact removal, despite these spikes reflecting physiological spike correlation. This excess removal of spike coincidences at *h*_0_ influences the effect of the correction: while the corrected surprise distribution (green) closely matches the original distribution (black) for large jitter widths (say, those larger than 1 ms), it diverges progressively from the original distribution for smaller jitter widths. Thus, this analysis reveals a limitation of our correction approach. It cannot compensate for the excess removal of extremely precise coincidences of correlated spikes. However, the correction works reasonably well for a wide range of coincidence precision, up to coincidences within ∼ 2 ms, which seems physiologically plausible. Smaller jitter widths (*<* 1ms) were studied to make the effect of wrongly removing precise coincidences on *h*_0_ more obvious.

Finally, we confirm if the correction works in the case where the correlated data also contain artifacts. Figure 6b shows the same graphs as in a, but obtained from correlated data also containing artifacts. Whereas the surprise distributions before artifact removal (black) largely shift towards right due to the coincidences that originate from the artifacts, the distribution after artifact removal (gray) closely matches between the data with and without artifacts, so is also the case for the surprise distribution after the correction (dashed green), for both wider (1 ms, left in each panel) and shorter (0.25 ms, right) jitter widths. The same observation holds for other jitter widths, as shown in Figure 6d. Despite the extremely large surprises in the data before artifact removal (black), the surprise distributions after artifact removal (gray) and further correction (green) are almost identical to those obtained from the data without artifacts (see Figure 6a,c). All these results confirm that the correction indeed works in the case of the UE_pop_ on the data containing artifacts, retaining as good performance as on the data without artifacts.

## 4 Discussion

In this study we reported that the removal of artifacts, identified in massively parallel spike trains as spike coincidences at the data sampling rate precision, has an impact on spike correlation analysis performed at a much lower time resolution. We focused here on the simplest and the most straightforward way of artifact removal, i.e., removal of all coincident spikes on the data sampling rate time scale (1/30 ms in popular setups). We demonstrated by use of simulated data and application of the UE_pop_ analysis that the artifact removal leads to an underestimation of the significance of the amount of spike coincidences in independent spike trains. We revealed that this is caused by an overestimation of the expected number of coincidences that is part of the UE_pop_ analysis, which needs to take into account the amount of the chance spike coincidences that occur on the data sampling time scale and are removed by the artifact removal. We here proposed a correction of the expected coincidence count for this effect, and demonstrated that this correction works for independent spike trains almost perfectly, and for correlated spike trains with certain limitations, no matter whether the data contain artifacts or not.

We found that the performance of the UE_pop_, in detecting excess spike synchrony in correlated spike trains, is degraded by the artifact removal even after the proposed correction, when the temporal precision of the coincidences of correlated spikes is set to be very high, e.g. more precise than ± 1 ms (Figure 6c). This happens because the artifact removal does not discriminate between artifactual and physiological coincidences but blindly removes all exact coincidences on the data sampling time resolution, and hence the removed coincidences contain more physiological coincidences as their precision becomes higher. The proposed correction cannot compensate for this excess removal of physiological coincidences, because it is based solely on the estimate of chance coincidences (on the sampling rate time scale), assuming no physiological correlations. In practice, however, this limitation would not cause serious problems when the UE_pop_ is applied to experimental spike recordings, because the biophysical constraints on the temporal precision of spiking activity of in vivo neurons is on the order of 1 or more ms (Riehle et al. (2000); Zandvakili and Kohn (2015)). Our analysis results indicate that, on this physiological temporal scale of coincidence precision, the performance of the UE_pop_ on data after artifact removal does provide a reasonable correction. The fact that the correlation for jitter widths smaller than 0.5ms further increases has another reason. UE_pop_ relies on binned and clipped (0-1) data. We apply disjunct binning, i.e., we do not have overlapping bins. This in turn may lead to a fission of synchronous events, e.g., for the case of a jitter of Δ = ± 1 ms and 1ms binning, there is a high chance that coincidences are distributed into 2 bins and thus not detected (Grün et al. (1999)). The smaller the jitter, the larger the probability that part of those coincidences fall into the 1ms bin and thus more are detected. This leads to a higher surprise. Thus, the results shown in Figure 6 contain a mix of effects: the smaller the jitter, the larger the amount of coincidences are detected, and on the other hand the more coincidences on the sampling rate time scale do exist and are removed as artifacts. This results with increasing jitter first in an increase of the corrected surprise and then to a decrease.

In Oberste-Frielinghaus et al. (2025), we reported that data sets of different labs and different recording techniques contain synchronous artifacts ranging across channels in the high-pass filtered raw recordings at the data sampling rate resolution, and we proposed a whitening technique to remove these artifacts based on the raw unfiltered recordings. However, often raw data are not available with publicly available spike data, which obviously have experienced preprocessing steps already, at least filtering and spike sorting. Then one would rather tend to remove the synchronous artifact events as we did in this study, in order to make sure that they are cleanly removed. We showed here that this may severely impact the subsequent analyses, in particular if fine temporal correlations in spike times are of interest. The removal of the artifacts and — by mistake, since they cannot be distinguished — chance events at the sampling resolution in turn reduces the significance of spike synchrony at a coarser time resolution and may lead to undetected excess spike synchrony, since the significance can be massively reduced. For the particular removal of the artifacts discussed here, we presented a correction that can be applied to mitigate this negative effect.

In Oberste-Frielinghaus et al. (2025), we also suggested to check if artifacts are in the data by computing the complexity distribution, based on the population time histogram of all spike trains at the sampling rate resolution, which illustrates how many synchronous events exist across the population. The complexity distribution, can be compared to that of an independent version of the data obtained by a surrogate generation method, e.g. by shifting the spike trains in time against each other up to a certain amount randomly (shift-dithering, see Stella et al. (2022)). This comparison is helpful to observe how many synchronous artifact events are in the data (see Oberste-Frielinghaus et al. (2025), Fig 1d). Unfortunately, the artifacts are not only above a certain complexity level, but are also mixed with uncorrelated chance synchronous events at lower complexities. Therefore, it is difficult to remove only the artifact events based solely on the complexity, though the complexity distribution is still a convenient tool to judge if a dataset is contaminated by synchronous artifacts or not.

One procedure to remove the artifacts in the raw data at a high time resolution is by whitening the data by using the zero-phase component analysis (ZCA) method by Bell and Sejnowski (1997). This approach assumes that the cross-talk across the recording channels is linear. In such a case this method removes only the artifacts, keeping the chance coincidences at the sampling rate time resolution intact, such that it should not affect the following spike correlation analysis on a longer time scale. The whitening step is often already an integrated part of the spike sorting method (e.g. in Kilosort (Pachitariu et al. (2024)) or Mountainsort (Chung et al. (2017))). Often there is a default setting in the sorting methods for how many of the recorded channels are included in one whitening step, typically smaller than the total number of parallel channels for reducing memory demand. How many channels are included in the whitening analysis is typically not obvious or mentioned in the metadata associated with publicly available datasets. However, this is an important piece of information, since we observed that, if the whitening does not include all channels but subsets of them, another artifact may be induced, e.g. electrode distance dependent artificial correlations. To be on the safe side, one should apply the whitening always to all channels simultaneously (Oberste-Frielinghaus et al. (2025)).

It turned out in this study that the UE_pop_ is a very sensitive detector for removed chance coincidence events on the recording time resolution. If the surprise values resulting from the UE_pop_ analysis are not distributed around 0, or above in case of the existence of excess spike synchrony, but rather below, one should be careful about potential impact of preprocessing. We assume that this effect is noticeable in any spike correlation method that compares to what is expected based on the firing rates, e.g. correlation coefficient etc. Likely the problem may not be noticed at all if the analysis is only based on firing rates.

## 5 Acknowledgements

We thank Dr. Alexander Thiele for useful discussions and Dr. Stefan Rotter for discussions in a previous project on that matter. Partly funded by the Deutsche Forschungsgemeinschaft (DFG, German Research Foundation) - grant 368482240/GRK2416 and grant 561027837/GR 1753/9-1, by the Ministry of Culture and Science of the State of North Rhine-Westphalia, Germany, and by the European Union’s Horizon Europe Programme under the Specific Grant Agreement No. 101147319 (EBRAINS 2.0 Project).

## Footnotes

1 Coincidentally this demonstrates that the UE_pop_is calibrated correctly: if the parallel spike trains follow a independent Poisson process the resulting coincidence statistic exactly matches what is statistically expected.

2 The given equations have another set of solutions, which have a positive sign for the square root term, but they represent unrealistically high firing probabilities and hence can be obviously rejected.

## References

Bedenbaugh P, Gerstein GL (1997) Multiunit normalized cross correlation differs from the average single-unit normalized correlation. Neural Comput 9:1265–1275

Bell AJ, Sejnowski TJ (1997) The “independent components” of natural scenes are edge filters. Vision research 37:3327–3338

Brochier T, Zehl L, Hao Y, Duret M, Sprenger J, Denker M, Grün S, Riehle A (2018) Massively parallel recordings in macaque motor cortex during an instructed delayed reach-to-grasp task. Sci Data 5:180055. 10.1038/sdata.2018.55

Chen X, Morales-Gregorio A, Sprenger J, Kleinjohann A, Sridhar S, van Albada SJ, Grün S, Roelfsema PR (2022) 1024-channel electrophysiological recordings in macaque V1 and V4 during resting state. Sci Data 9:77. 10.1038/s41597-022-01180-1

Chung JE, Magland JF, Barnett AH, Tolosa VM, Tooker AC, Lee KY, Shah KG, Felix SH, Frank LM, Greengard LF (2017) A fully automated approach to spike sorting. Neuron 95:1381–1394.e6. 10.1016/j.neuron.2017.08.030

Grün S, Abeles M, Diesmann M (2008) Impact of higher-order correlations on coincidence distributions of massively parallel data. In: Lecture Notes in Computer Science, ‘Dynamic Brain - from Neural Spikes to Behaviors’, vol 5286

Grün S, Diesmann M, Aertsen A (2002a) Unitary events in multiple single-neuron spiking activity: I. detection and significance. Neural Comput 14:43–80

Grün S, Diesmann M, Aertsen A (2002b) Unitary events in multiple single-neuron spiking activity: II. nonstationary data. Neural Comput 14:81–119

Grün S, Diesmann M, Grammont F, Riehle A, Aertsen A (1999) Detecting unitary events without discretization of time. J Neurosci Methods 94:67–79

Louis S, Gerstein GL, Grün S, Diesmann M (2010) Surrogate spike train generation through dithering in operational time. Front Comput Neurosci 4:127

Maldonado P, Babul C, Singer W, Rodriguez E, Berger D, Grün S (2008) Synchronization of neuronal responses in primary visual cortex of monkeys viewing natural images. J Neurophysiol 100:1523–1532

Oberste-Frielinghaus J, Morales-Gregorio A, Essink S, Kleinjohann A, Barthélemy F, Musall S, Grün S, Ito J (2025) Detection and removal of hyper-synchronous artifacts in massively parallel spike recordings. BioRxiv 10.1101/2024.01.11.575181, v2

Pachitariu M, Sridhar S, Pennington J, Stringer C (2024) Spike sorting with Kilosort4. Nat Methods 21:914–921. 10.1038/s41592-024-02232-7, Publisher: Nature Publishing Group

Palm G (1981) Evidence, information and surprise. Biol Cybern 42:57–68

Pazienti A, Grün S (2006) Robustness of the significance of spike synchrony with respect to sorting errors. J Comput Neurosci 21:329–342

Pei F, Ye J, Zoltowski DM, Wu A, Chowdhury RH, Sohn H, O’Doherty JE, Shenoy KV, Kaufman MT, Churchland M, Jazayeri M, Miller LE, Pillow J, Park IM, Dyer EL, Pandarinath C (2021) Neural latents benchmark ‘21: Evaluating latent variable models of neural population activity. In: Advances in Neural Information Processing Systems (NeurIPS), Track on Datasets and Benchmarks

Riehle A, Grammont F, Diesmann M, Grün S (2000) Dynamical changes and temporal precision of synchronized spiking activity in monkey motor cortex during movement preparation. J Physiol 94:569–582

Staude B, Rotter S, Grün S (2010) CuBIC: cumulant based inference of higher-order correlations in massively parallel spike trains. J Comput Neurosci 29:327–350. 10.1007/s10827-009-0195-x

Steinmetz N, Carandini M, Harris KD (2019) “Single Phase3” and “Dual Phase3” Neuropixels Datasets

Stella A, Bouss P, Palm G, Grün S (2022) Comparing surrogates to evaluate precisely timed higher-order spike correlations. eNeuro 9:ENEURO.0505–21.2022. 10.1523/ENEURO.0505-21.2022

Torre E, Quaglio P, Denker M, Brochier T, Riehle A, Grün S (2016) Synchronous spike patterns in macaque motor cortex during an instructed-delay reach-to-grasp task. J Neurosci 36:8329–8340. 10.1523/jneurosci.4375-15.2016

Yu BM, Cunningham JP, Santhanam G, Ryu SI, Shenoy KV, Sahani M (2009) Gaussian-Process Factor Analysis for Low-Dimensional Single-Trial Analysis of Neural Population Activity. J Neurophysiol 102:614–635. 10.1152/jn.90941.2008

Zandvakili A, Kohn A (2015) Coordinated neuronal activity enhances corticocortical communication. Neuron 87:827–839. 10.1016/j.neuron.2015.07.026

